# Dynamic constraints on activity and connectivity during the learning of value

**DOI:** 10.1101/448464

**Authors:** Azeez Adebimpe, Maxwell Bertolero, Ankit N. Khambhati, Marcelo G. Mattar, Daniel Romer, Sharon L. Thompson-Schill, Danielle S. Bassett

## Abstract

Human learning is a complex process in which future behavior is altered via the modulation of neural activity. Yet, the degree to which brain activity and functional connectivity during learning is constrained across subjects, for example by conserved anatomy and physiology or by the nature of the task, remains unknown. Here, we measured brain activity and functional connectivity in a longitudinal experiment in which healthy adult human participants learned the values of novel objects over the course of four days. We assessed the presence of constraints on activity and functional connectivity using an inter-subject correlation approach. Constraints on activity and connectivity were greater in magnitude than expected in a non-parametric permutation-based null model, particularly in primary sensory and motor systems, as well as in regions associated with the learning of value. Notably, inter-subject connectivity in activity and connectivity displayed marked temporal variations, with inter-subject correlations in activity exceeding those in connectivity during early learning and *visa versa* in later learning. Finally, individual differences in performance accuracy tracked the degree to which a subject’s connectivity, but not activity, tracked subject-general patterns. Taken together, our results support the notion that brain activity and connectivity are constrained across subjects in early learning, with constraints on activity, but not connectivity, decreasing in later learning.

## Introduction

Each human brain is unique, but all brains share a common form. Functional connectivity, which measures the statistical similarity between activity time series of brain regions, can vary appreciably across individuals, and can serve as an individual’s fingerprint [1]. Moreover, recent work has demonstrated that functional connectivity can predict task performance [2] and the capacity for skill learning [3, 4, 5]. Similarly, a single cognitively demanding task can elicit quite different patterns and magnitudes of activity in different individuals [6]. Finally, individual differences in functional connectivity during rest are related to individual differences in brain activity during task performance [7]. Despite these notable instances of variance in brain activity and connectivity, there also exist several subject-general constraints; macro-organizational principles of functional connectivity appear to be conserved throughout healthy normative populations [8, 9], and there are reliable group level task-induced patterns of brain activity [10].

Similarly, learning is a dynamic process with both variance and conservation across individuals; everyone learns differently, but much of the mechanics of learning follow a common form. While individual differences in brain and behavior certainly exist during learning, two notable factors constrain neural processes to be markedly similar across individuals. First, engagement in the same task can constrain brain activity and functional connectivity. The mechanics of the particular skill that is learned, the information that one learns, and learning strategies, if similar across subjects, can induce similarities in neural dynamics. Second, conserved functional and structural architecture can constrain brain activity and connectivity [11]. Unique and shared genetic and environmental factors present in the early stages of development can give rise to distinct and common wiring patterns, respectively [12], which can in turn lead to distinct and conserved spatiotemporal responses in neuronal ensembles [13, 14, 15]. The two factors of task engagement and brain wiring can also interact with one another: two subjects with similar brain wiring that are employing the same strategy to learn the same skill will likely exhibit more similar brain activity and connectivity patterns than two individuals with different brain wiring that are employing different learning strategies.

However, the nature of constraints on activity and connectivity during learning, due to task engagement or similar brain architectures, is unknown. The regional variance in constraints is also not known; in other words, we do not know whether some brain regions are more or less constrained than others. Furthermore, we do not know whether some regions’ activity is more constrained that their connectivity, or *vice versa*. Moreover, the temporal variance in constraints on activity and connectivity, and any relationship between the two, during learning is unknown. What is the temporal and dynamical evolution of constraints on activity and connectivity during learning? Can constraints on activity and functional connectivity distinguish between different phases of value learning? Finally, how do constraints on activity and connectivity relate to learning performance?

Here we use intersubject connectivity (ISC) and intersubject functional connectivity (ISFC) to assess the extent to which each region’s activity and connectivity are constrained across subjects during the course of value learning. We test three specific hypotheses. First, we hypothesize that the strongest constraints would exist primarily in sensory and motor systems, as well as in regions associated with the learning of value, as these regions are either constrained across subjects in general or are constrained because of similar learning processes. Second, we hypothesize that the coherence of stimulus-induced activity would place constraints on brain activity and connectivity during early learning, but that subject-specific activity and connectivity patterns would dominate later learning when greater neural real estate was available for non-task-related processing as well as processing reflecting subject-specific learning strategies. Third and finally, we hypothesize that the extent to which a subject’s brain activity or connectivity obeys subject-general constraints relates to learning performance. Altogether, our investigations demonstrate the presence of dynamic constraints on task-dependent activity and functional connectivity during the learning of value.

## Materials and Methods

### Experimental setup and procedure

Participants learned the monetary value of 12 novel visual stimuli in a reinforcement learning paradigm. Learning occurred over the course of four MRI scan sessions conducted on four consecutive days. The novel stimuli were 3-dimensional shapes generated with a custom built MATLAB toolbox. Publicly available code is located here: http://github.com/saarela/ShapeToolbox. ShapeToolbox allows the generation of three-dimensional radial frequency patterns by modulating basis shapes, such as spheres, with an arbitrary combination of sinusoidal modulations in different frequencies, phases, amplitudes, and orientations. A large number of shapes were generated by selecting combinations of parameters at random. From this set, we selected twelve that were considered to be sufficiently distinct from one another. A different monetary value, varying from $1.00 to $12.00 in integer steps, was assigned to each shape. These values were not correlated with any parameter of the sinusoidal modulations, so that visual features were not informative of value.

Participants completed approximately 20 minutes of the main task protocol on each scan session, learning the values of the 12 shapes through feedback. The sessions were comprised of three scans of 6.6 minutes each, starting with 16.5 seconds of a blank gray screen, followed by 132 experimental trials (2.75 seconds each), and ending with another period of 16.5 seconds of a blank gray screen. Stimuli were back-projected onto a screen viewed by the participant through a mirror mounted on the head coil and subtended 4 degrees of visual angle, with 10 degrees separating the center of the two shapes. Each presentation lasted 2.5 seconds and, at any point within a trial, participants entered their responses on a 4-button response pad indicating their shape selection with a leftmost or rightmost button press. Stimuli were presented in a pseudorandom sequence with every pair of shapes presented once per scan.

Feedback was provided as soon as a response was entered and lasted until the end of the stimulus presentation period. Participants were randomly assigned to two groups depending on the type of feedback received. In the RELATIVE feedback case, the selected shape was highlighted with a green or red square, indicating whether the selected shape was the most valuable of the pair or not, respectively. In the ABSOLUTE feedback case, the actual value of the selected shape (with variation) was displayed in white font. After each run, both groups received feedback about the total amount of money accumulated in the experiment up to that point. The experimental protocol has been reported previously [4].

### MRI data collection and preprocessing

Magnetic resonance images were obtained at the Hospital of the University of Pennsylvania using a 3.0 T Siemens Trio MRI scanner equipped with a 32-channel head coil. T1-weighted structural images of the whole brain were acquired on the first scan session using a three-dimensional magnetization-prepared rapid acquisition gradient echo pulse sequence with the following parameters: repetition time (TR) 1620 ms, echo time (TE) 3.09 ms, inversion time 950 ms, voxel size 1 mm by 1 mm by 1 mm, and matrix size 190 by 263 by 165. To correct geometric distortion caused by magnetic field inhomogeneity, we also acquired a field map at each scan session with the following parameters: TR 1200 ms, TE1 4.06 ms, TE2 6.52 ms, flip angle 60^°^, voxel size 3.4 mm by 3.4 mm by 4.0 mm, field of view 220 mm, and matrix size 64 by 64 by 52. In all experimental runs with a behavioral task, T2*-weighted images sensitive to blood oxygenation level-dependent contrasts were acquired using a slice accelerated multi-band echo planar pulse sequence with the following parameters: TR 2000 ms, TE 25 ms, flip angle 60^°^, voxel size 1.5 mm by 1.5 mm by 1.5 mm, field of view 192 mm, and matrix size 128 by 128 by 80. In all resting state runs, T2*-weighted images sensitive to blood oxygenation level-dependent contrasts were acquired using a slice accelerated multi-band echo planar pulse sequence with the following parameters: TR 500 ms, TE 30 ms, flip angle 30^°^, voxel size 3.0 mm by 3.0 mm by 3.0 mm, field of view 192 mm, matrix size 64 by 64 by 48.

Cortical reconstruction and volumetric segmentation of the structural data was performed with the Freesurfer image analysis suite [16]. Boundary-Based Registration between structural and mean functional image was performed with Freesurfer bbregister [17]. Preprocessing of the fMRI data was car-ried out using FEAT (FMRI Expert Analysis Tool) Version 6.00, part of FSL (FMRIB’s Software Library, www.fmrib.ox.ac.uk/fsl). The following pre-statistics processing was applied: EPI distortion correction using FUGUE [18], motion correction using MCFLIRT [19], slice-timing correction using Fourier-space time series phase-shifting, non-brain removal using BET [20], grand-mean intensity normalization of the entire 4D dataset by a single multiplicative factor, and highpass temporal filtering via Gaussian-weighted least-squares straight line fitting with *σ* = 50.0*s*. Nuisance time series were voxelwise regressed from the preprocessed data. Nuisance regressors included (i) three translation (X, Y, Z) and three rotation (pitch, yaw, roll) time series derived by retrospective head motion correction (R = [X, Y, Z, pitch, yaw, roll]), together with the first derivative and square expansion terms, for a total of 24 motion regressors [21]); (ii) the first five principal components of non-neural sources of noise, estimated by averaging signals within white matter and cerebrospinal fluid masks, obtained with Freesurfer segmentation tools, and removed using the anatomical CompCor method (aCompCor) [22]; and (iii) a measure of a local source of noise, estimated by averaging signals derived from the white matter region located within a 15 mm radius from each voxel, using the ANATICOR method [23]. Global signal was not regressed out of voxel time series [24, 25].

We parcellated the brain into 112 cortical and subcortical regions, separated by hemisphere using the structural Harvard-Oxford atlas of the FMRIB (Oxford Centre for Functional Magnetic Resonance Imaging of the Brain) Software Library (FSL; Version 5.0.4) [26, 27]. We warped the MNI152 regions into subject-specific native space using FSL FNIRT and nearest-neighbor interpolation and calculated the average BOLD signal across all gray matter voxels within each region. The participant’s gray matter voxels were defined using the anatomical segmentation provided by Freesurfer, projected into the subject’s EPI space with *bbregister*. For each individual scan, we extracted regional mean BOLD time series by averaging voxel time series in each of the 112 regions of interest.

### Brain network construction

To perform network analyses one must define the two most fundamental elements of the network – nodes and edges. These two elements are the building blocks of networks and their accurate definitions are very important for any network models [28]. The standard method of defining network nodes in the field of network neuroscience is to consider neuroimaging data such as fMRI and apply a structural atlas or parcellation that separates the whole brain volume into different regions defined by known anatomical differences [29]. A network node thus represents the collection of voxels within a single anatomically defined region. A network edge reflects the statistical dependency between the activity time series of two nodes. In this study, the brain is parcellated into 112 subcortical and cortical regions defined by the structural Harvard-Oxford atlas of the fMRIB [26, 27]. Each region’s activity is given by the mean time series across all voxels within that region.

The edge weights that link network nodes were given by the wavelet transform coherence (WTC) [30], smoothed over time and frequency to avoid bias toward unity coherence. Specifically, we use Morlet wavelets with coefficients given by:

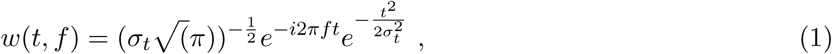

where *f* is the center frequency and *σ_t_* is the temporal standard deviation. The time-frequency estimate, *X*(*t,f*) of time series *x*(*t*) was computed by a convolution with the wavelet coefficients:

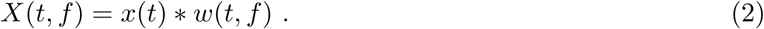

We selected the central frequency of 1/12 Hz corresponding to a spectral width of 0.05 to 0.11 Hz for full width at half maximum. Then the wavelet transform coherence between two time series *x(t)* and *y(t)* is defined as follows [30, 31, 32]:

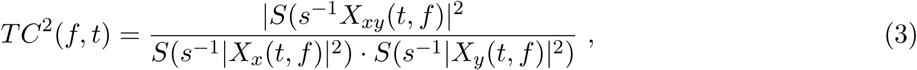

where *X_xy_* is the cross-wavelet of *X_x_* and *X_y_*, *s* is the scale which depends on the frequency [31, 32], and *S* is the smoothing operator. This definition closely resembles that of a traditional coherence, with the marked difference that the wavelet coherence provides a localized correlation coefficient in both time and frequency. Higher scales are required for lower frequency signals and in this study, we used *s* = 32 for the smoothing operation. This procedure was repeated for all pair of regions yielding the 112 by 112 adjacency matrix, A, representing the functional connectivity between brain regions.

### Network modularity

In network neuroscience, the term modularity can be used to refer to the concept that brain regions cluster into modules or *communities* [33]. These communities can be identified computationally using machine learning techniques in the form of community detection algorithms [34]. A community of nodes is a group of nodes that are tightly interconnected. In this study, we implemented a generalized Louvain-like community detection algorithm [35, 36] that considers multiple adjacency matrices as slices of a multilayer network, and which then produces a partition of brain regions into modules that reflects each subject’s community structure across the multiple stages of learning instantiated in the four days of task practice. The multilayer network was constructed by connecting the adjacency matrices of all scans and subjects with interlayer links. We then maximized a multilayer modularity quality function, Q, that seeks a partition of nodes into communities in a way that maximizes intra-community connections [36]:

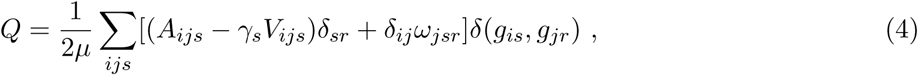

where *A_ijs_* is the *ij*^th^ element of the adjacency matrix of slice *s*, and element *V_ijs_* is the component of the null model matrix tuned by the structural resolution parameter *γ*. In this study, we set *γ* = 1, which is the standard practice in the field when no *a priori* hypotheses exist to otherwise inform the choice of *γ*. We employed the Newman-Girvan null model within each layer by using 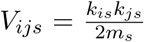, where *k* is the total edge weight and *m_s_* is the total edge weight in slice *s*. The interslice coupling parameter, *ω_jsr_*, is the connection strength of the interlayer link between node *j* in slice *s* and node *j* in slice *r,* and the total edge weight in the network is 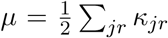. The node strength, *κ_jr_*, is the sum of the intraslice strength and interslice strength: *κ_jr_* = *κ_jr_* + *c_jr_*, and *c_jr_* = *Σ_s_ω_jrs_*. In this study, we set *ω* = 1, which is the standard practice in the field when no *a priori* hypotheses exist to otherwise inform the choice of *ω*. Finally, the indicator δ(*g_i_*, *g_j_*) = 1 if nodes *i* and *j* are assigned to the same community, and is 0 otherwise.

We obtained a partition of the brain into communities for each scan and subject. To obtain a single representative partition of brain regions into distinct communities, we computed a module allegiance matrix [37, 38, 39], whose *ij*^th^ entry represents the probability that region *i* and region *j* belong to the same community across scans and participants. We then applied single-layer modularity maximization to this module allegiance matrix to obtain a single partition of brain regions into consensus modules [38]. The seven network communities generated with this procedure are shown in Fig. 5.

### Edge strength

In complementary analyses, we also investigated which regions of the brain were characterized by high strength within the network. The edge strength of node *i* is defined as

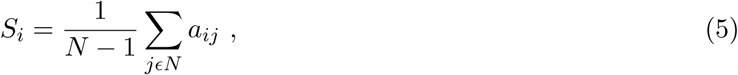

where *α_ij_* is the *ij*^th^ element of the adjacency matrix with *N* nodes.

### Statistical Methods: Inter-subject correlation (ISC) and inter-subject functional connectivity(ISFC)

We used the Pearson correlation coefficient to compute the inter-subject correlation (ISC) of each brain region for each subject. First, we calculated the region-wise temporal correlation between every pair of subjects as:

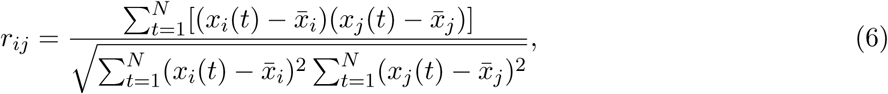

where *N* is the number of points in the time series data, and where *r_ij_* is the correlation coefficient of a region between the times series *x_i_* and *x_j_* of the *i*th and *j*th subjects, respectively.

To test the statistical significance of the correlation between the fMRI BOLD signals of a single region from two subjects, we performed a fully non-parametric permutation test with 10,000 randomizations [40]. This test accounts for slow-scale autocorrelation structure in the BOLD time series [41] by removing phase information from each BOLD signal through Fourier phase randomization. We repeated this procedure 10,000 times to obtain a null distribution of the maximum noise correlation values, and we defined the threshold for a pair correlation as the *q* × 100^*th*^ percentile of the null distribution of maximum values.

To obtain the ISC, we averaged only significant correlation values out of 190 correlation values, *r_ij_* from all subject pairs, to obtain one ISC for each region:

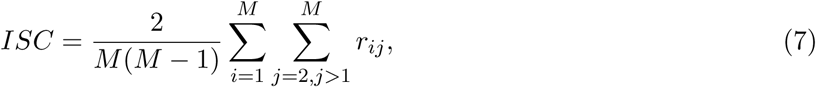

where *M* is the number of subjects. This same procedure was followed for each scan, for each day, and for both rest and task conditions.

Inter-subject functional connectivity (ISFC) was obtained from the functional connectivity (FC) matrices of all of the subjects, and can be thought of as an estimate of the correlation in FC between one subject and all other subjects. Specifically, we computed the ISFC of each subject as the correlation between the single subject FC matrix and the average of all other subject-specific FC matrices as

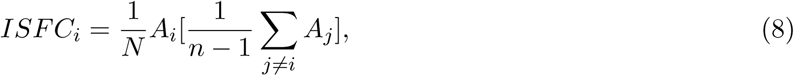

where **A** is the functional connectivity matrix of a subject, and where *n* and *N* are the total number of subjects and total number of regions, respectively.

### Methodological Considerations

The number of subjects analyzed here is smaller than large-scale data collections [42]. However, given the four day experimental design involving the same subjects, this dataset nevertheless represents a rich opportunity to analyze the temporal evolution of constraints on activity and connectivity during learning. We hope that future large-scale data collections will include similar experiments. We chose a simple and intuitive method to measure the similarity of brain activity and connectivity across subjects, but certainly more sophisticated methods could be developed and will likely uncover results that are complimentary to the results reported here. Finally, there are many ways to parcellate the brain into regions, represented by nodes in network analyses. While many parcellations maximize the similarity of functional connectivity within each parcel [43], given that we analyzed both activity and connectivity, and brain activity often crosses boundaries defined by these functional parcellations, we chose an anatomical parcellation with relatively large parcels so as not to bias our results towards activity or connectivity. Moreover, this atlas has been used extensively in prior literature.

## Results

We investigated functional mechanisms that facilitate network reorganization associated with value learning by assessing ISC (Fig. 1a) and ISFC (Fig. 1b) estimated from functional MRI (fMRI) data acquired in 20 healthy subjects (9 females; ages 19–53 years; mean age = 26.7 years) over four consecutive days. Each scanning session contained both data collected while the subject rested and data collected while the subject engaged in a value-learning task. From each session, we extracted the BOLD time series of 112 cortical and subcortical regions defined by the Harvard-Oxford atlas [26, 27]. We measured ISC by estimating the correlation between regional BOLD signal time series for each pair of *S* subjects. This procedure resulted in an *S* × *S* correlation matrix for each brain region. Next, we constructed a functional connectivity matrix for each subject and each scan, where each *ij*^th^ element in the *N* × *N* matrix indicated the wavelet coherence between the time series of region *i* and the time series of region *j*. We then measured ISFC for a given subject and region *i* by computing the Pearson correlation between row *i* in that subject’s functional connectivity matrix and the average of row *i* in all of the other subjects’ functional connectivity matrices. We performed this calculation for every region, resulting in an ISFC array of length *N* for each subject (see Materials and Methods).

**Figure 1.**
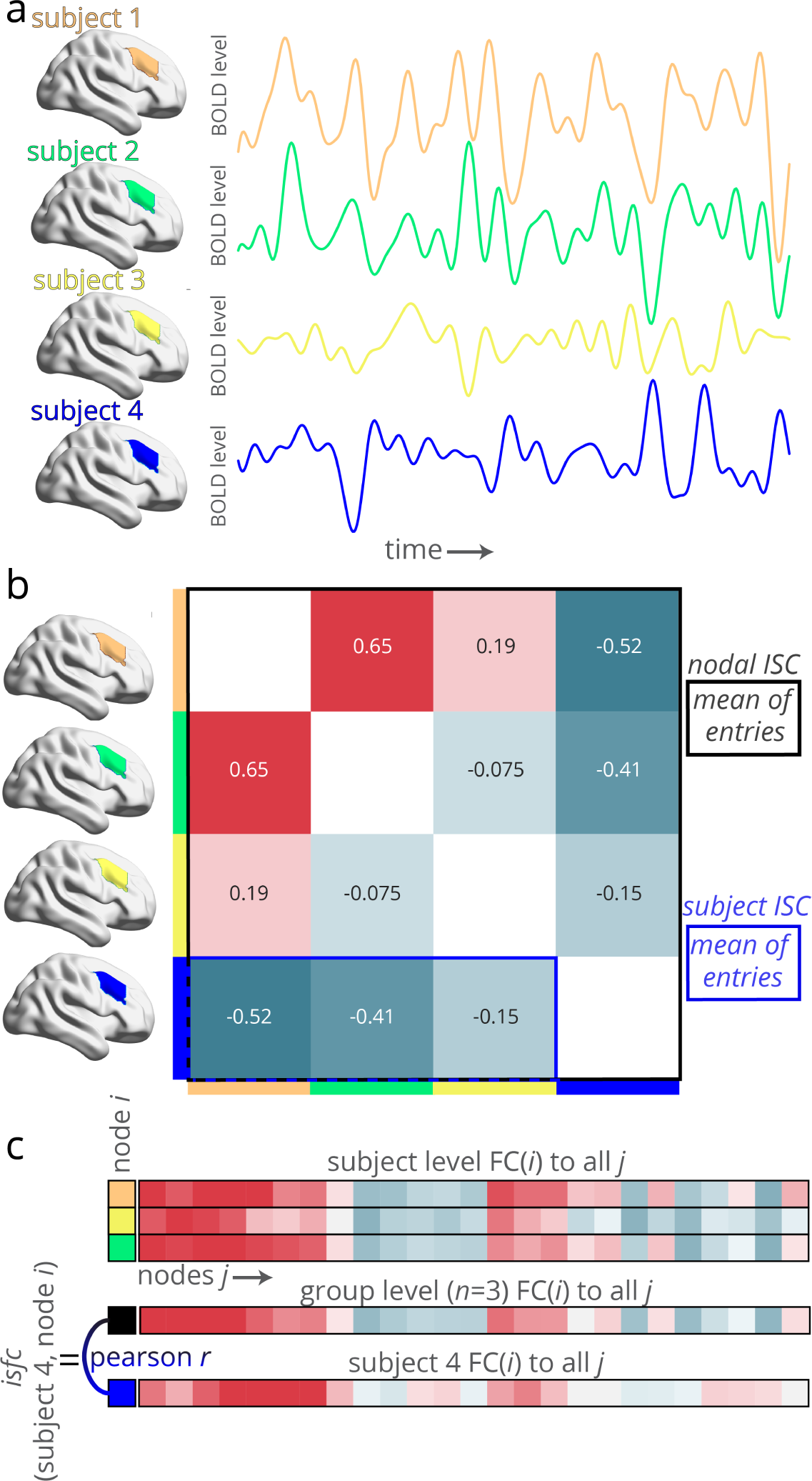
Measuring intersubject connectivity and intersubject functional connectivity. **a**, To measure inter-subject connectivity (ISC), the mean time series of a single region (or network node) is extracted in each subject. Here we provide an illustrative depiction with four subjects. **b**, Next, the correlation between all pairs of subjects’ time series is calculated. The mean of all entries (black box) is the region’s ISC. The mean of a subject’s row (blue box) is the subject’s ISC. **c**, To measure inter-subject functional connectivity for a single subject (subject #4) and region *i*, the functional connectivity between region *i* and all other regions *j* is calculated for each subject. Next, the mean connectivity between region *i* and all other regions *j* is calculated without subject #4 (black box). The ISFC for subject #4 and region *i*, then, is the correlation between the mean connectivity between region *i* and all other regions *j* (without subject #4) and the connectivity between region *i* and all other regions *j* in subject #4.

### Constraints on activity and connectivity during learning

First, we asked whether constraints on brain activity and functional connectivity are homogeneous across brain areas, or whether some areas are more constrained than others. We hypothesized that the strongest constraints would exist primarily in sensory and motor systems, as well as in regions associated with the learning of value, as these regions are either constrained across subjects in general or are constrained because of similar learning processes. More specifically, we expected subject-general constraints in (i) motor cortex, consistent with the shared demands of finger movements necessary to press buttons on the response box [37, 44], (ii) visual cortex, consistent with the shared demands of cognitive processing necessary to parse the visual stimuli of the novel objects, and (iii) other areas previously associated with the learning of value, including orbital frontal cortex and lateral occipital cortex [45].

To test these hypotheses, we assessed which brain regions exhibited ISC values that were significantly greater than expected across task sessions (*t*-test, *p* < 0.05, corrected for multiple comparisons with 10000 random permutations [46]) across all task sessions. We observed significant ISC broadly across the cortex (Fig. 2a). Next, we assessed which brain regions exhibited ISFC values that were significantly greater than expected across sessions (*t*-test, *p* < 0.05, corrected for multiple comparisons). We also observed significant ISFC (*t*-test, *p* < 0.05 corrected for multiple comparisons) broadly across the cortex, and specifically in sensorimotor and visual regions (Fig. 2b). The spatial differences in ISC and ISFC across the cortex were investigated further by *z*-scoring the values of each and subtracting them from one another (Fig. 2c,d). While some differences existed between the brain regions displaying high ISC and the brain regions displaying high ISFC, we observed a positive correlation between regions’ ISFC and ISC (*r* = 0.48, *p* = 9.71 × 10^−8^).

**Figure 2.**
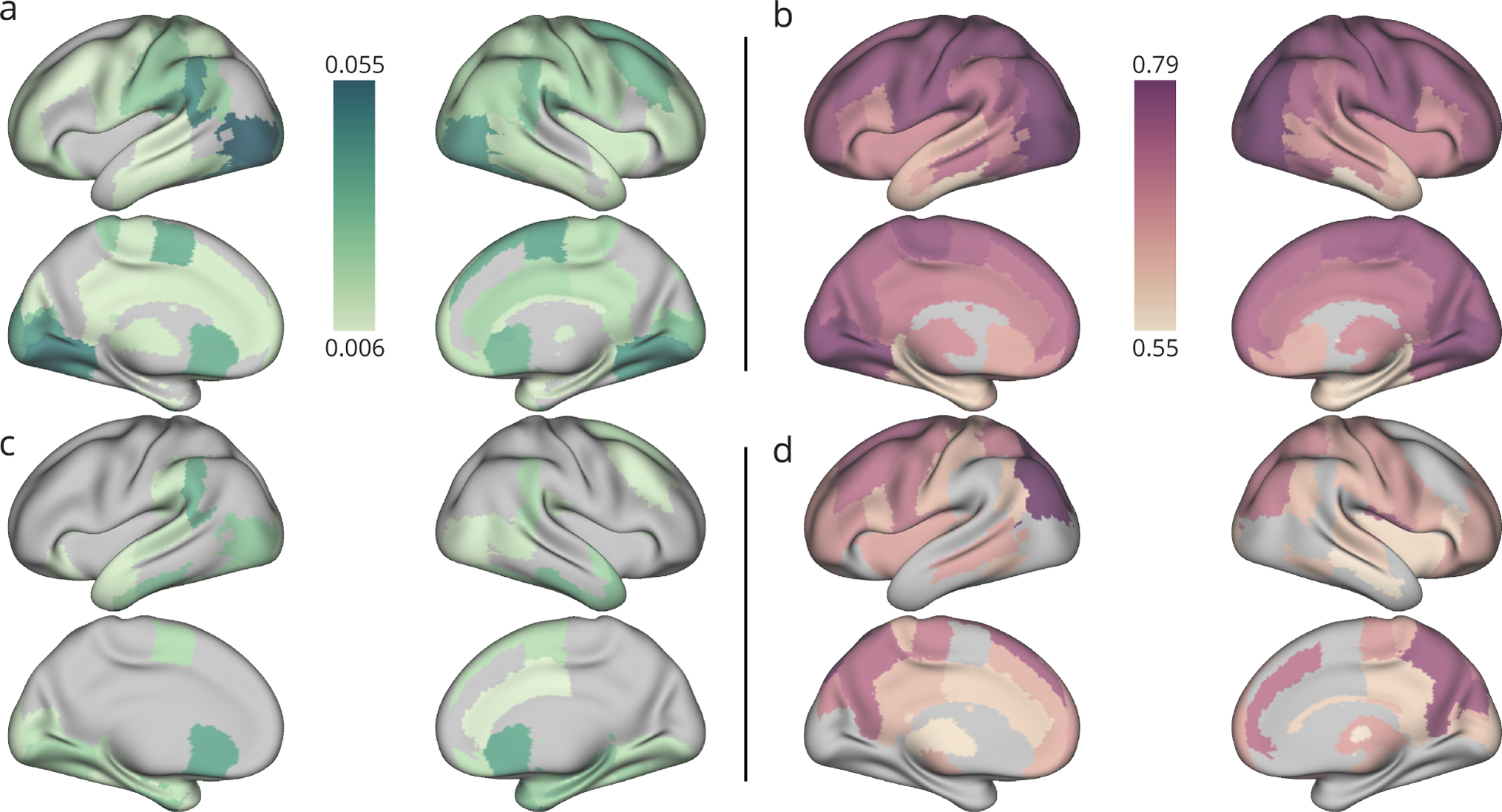
Intersubject connectivity and intersubject functional connectivity during learning. **a**, In the fMRI data acquired during the performance of the value learning task, we observed significant ISC (*p* < 0.05, corrected for multiple comparisons) in the lateral occipital cortex, lingual gyrus, supramarginal, sensorimotor areas, and anterior cingulate. **b**, Again in the fMRI data acquired during the performance of the value learning task, we observed significant ISFC (*p* < 0.05, corrected for multiple comparisons) in the pre-central, lingual gyrus, and left supramarginal, as well as in the precuneus, cuneus, frontal pole, and cingulate. **c**–**d**, To visualize the spatial differences in ISC and IFSC, the values were *z*-scored separately and then subtracted from one another.

While we observed high ISFC and ISC in motor and visual regions, to empirically determine which regions with high ISC or ISFC are associated with value, we used the “value” term association map in Neurosynth [47], which measures how often each voxel is reported in studies with “value” in the abstract (*n* = 470) versus all other studies. Fig. 3a displays this map, which primarily loads on to medial orbital frontal cortex, but also loads on to the canonical fronto-parietal network [48] regions. In order to find regions with high ISC or ISFC that are also associated with value tasks, we multiplied the *t*-values from the meta-analysis by the *z*-scored ISFC and ISC values, retaining the positive values (Fig. 3b,c). While ISC was high at regions canonically associated with value learning such as the orbital frontal cortex, we found that the ISFC was high in regions more loosely associated with learning in the fronto-parietal network areas.

**Figure 3.**
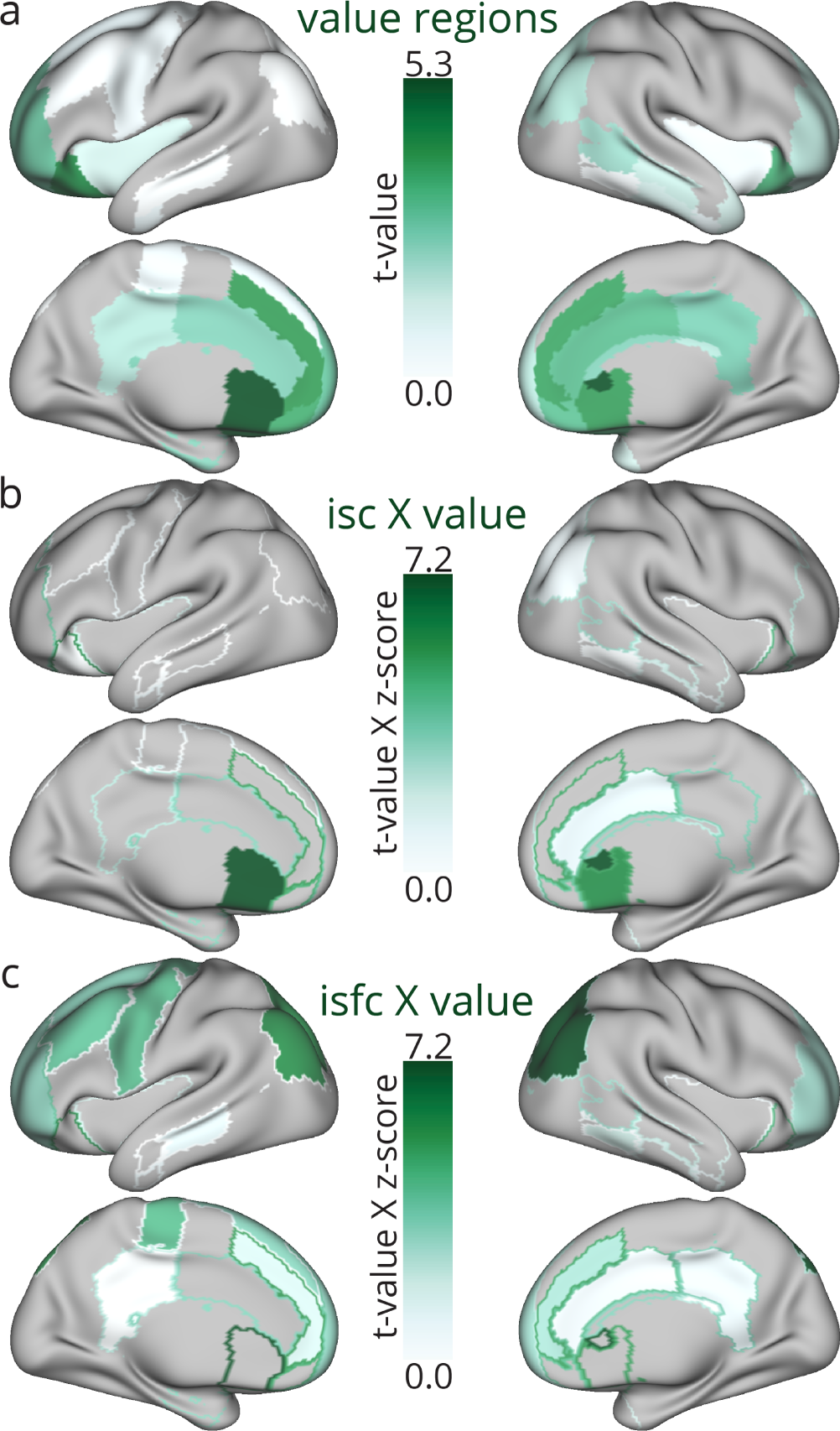
Constraints on activity and connectivity in regions associated with value. **a**, Regions that are associated with value tasks in NeuroSynth (*n* = 470) [47]. **b**, Regions that are associated with value and have high ISC. **c**, Regions that are associated with value and have high ISFC. In panels **b**–**c**, the values from panel a are shown in outlines.

Collectively, these results suggest that motor, visual, and value learning regions have high constraints on their activity and connectivity. While constraints appear to be mostly similar on activity and connectivity, it is possible that activity is more constrained in regions that encode or represent value, while connectivity is more constrained in regions associated with control processes or other learning-related processes.

### Dynamic constraints on functional architecture during learning

Next we asked whether constraints on activity and functional connectivity could distinguish between different phases of value learning. We expected that brain activity and connectivity would be more highly constrained across subjects during early learning, as indicated by high values of ISC and ISFC. This hypothesis is driven by the notion that, in the later stages of learning, which are characterized by high performance accuracy and increasing automaticity of responses, greater neural real estate might be available for contemporaneous, non-task-related processing as well as processing reflecting subject-specific learning strategies. However, in late learning, we still expected that functional connectivity would remain more heavily constrained than activity, due to its temporally extended nature being driven by commonly reinforced patterns of stimulus response, as well as due to its sensitivity to underlying brain anatomy conserved across subjects.

To test these hypotheses, we first examined the dynamics of ISC during value learning (Fig. 4). We found that the average ISC over all brain regions increased from the first day to the second day, and then subsequently decreased through the fourth day, with the decrease on the final day being significant (*t* = 3.21, *p* = 0.004; Fig. 4b). These trends in task-related ISC dynamics suggest that ISC might support two phases of value learning: (i) increasing ISC between day one and day two may be associated with increasing constraints on activity, perhaps as a result of common neurophysiological mechanisms across subjects that facilitate early stage learning of the task mechanics, and (ii) decreasing ISC from day two through day four may be associated with less constrained dynamics, perhaps as subjects explore diverse cognitive strategies to further increase their performance on the task. In a complementary analysis, we also examined the dynamics of ISFC during value learning, and found that ISFC also increased from day one to day two (*t* = 3.37, *p* = 0.003), peaking at day two (Fig. 4b), suggesting that activity and functional connectivity are both most constrained to common organizational rules across the group in early stages of learning.

**Figure 4.**
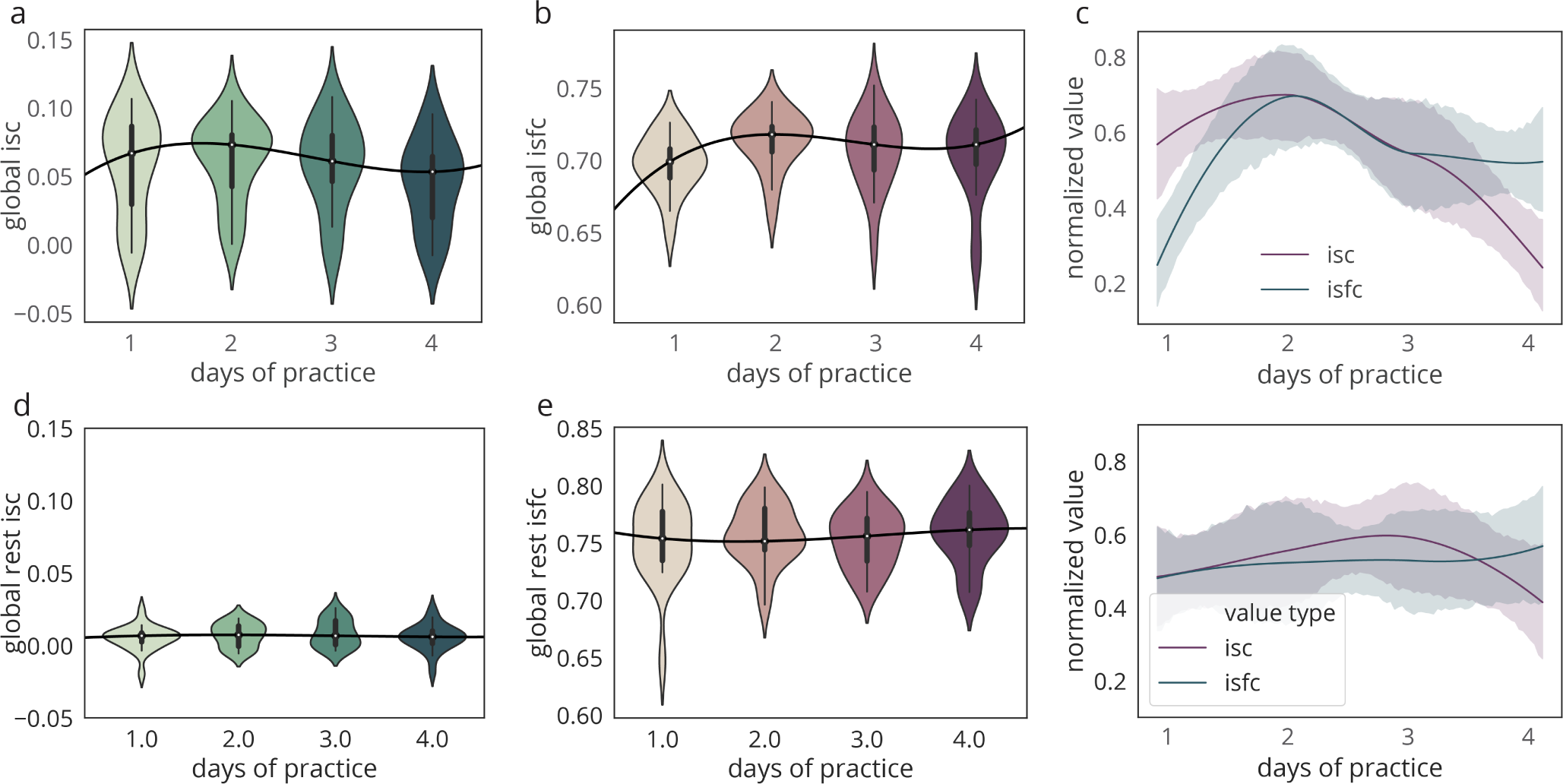
Temporal dynamics of whole-brain ISC and ISFC during value learning. **a** We observed that the ISC increased from day one to day two of task practice, and then decreased in day three and day four. **b** We observed that the ISFC increased from day one to day two of task practice, thereafter remaining relatively steady. **c** The temporal dynamics of the normalized ISC and ISFC, which both peak on day two; however, ISFC becomes significantly higher in day four. **d**–**f** Same as in panels **a**–**c**, but in this case the ISC and ISFC were measured during the resting state. Note that the temporal dynamics observed during the learning sessions were not observed during the resting state.

Next, we explicitly tested for potential interactions between ISC and ISFC over time. We found that ISC and ISFC were significantly positively correlated on each of the four days (day 1: *r* = 0.4647, *p* = 3.42 × 10^−7^, day 2: *r* = 0.4274, *p* = 3.26 × 10^−6^, day 3: *r* = 0.3946, *p* = 1.96 × 10^−5^, day 4: *r* = 0.4956, *p* = 4.27 × 10^−8^), suggesting that increased group-level constraints on activity are related to increased group-level constraints on functional connectivity. Nevertheless, we also observed some evidence for a divergence in the constraints on ISC and ISFC during the later stages of learning; for these comparisons, both ISC and ISFC were normalized by subtracting the minimum and dividing by the range. Specifically, on day four we observed that normalized ISFC values were significantly greater than normalized ISC values (Fig. 4c), suggesting that activity is more autonomous than functional connectivity.

As a null model, we computed ISC for the rest condition, during which fMRI data was also acquired in each of the four days of the experiment. We observed that the average ISC was much lower during the rest condition compared to during the value learning task – the average ISC was statistically indistinguishable from zero. This observation is perhaps not so surprising when one considers that during the rest condition, activity is no longer time locked to any stimulus, and therefore subject’s brain dynamics are allowed to evolve independently. Moreover, we observed that the ISC was similar across all four days of the resting condition, not varying appreciably across days (Fig. 4d). Similarly, we observed that the ISFC was similar across all four days of the resting condition, not varying appreciably across days (Fig. 4e). Thus, our results regarding the temporal evolution of ISC and ISFC are driven by learning, not time, as they are only observed during the learning condition, and not during the resting condition.

Finally, we verified that the average ISC estimated during task performance over all four days of training was significantly greater than the ISC estimated during the resting state over those same four days (*t* = 7.94, *p* = 1.87 × 10^−7^). Moreover, we observed no correlation between each subject’s global ISC during learning and during the resting state (*r*=0.10, *p*=0.66). While the manner in which brain regions similarly activate across subjects is certainly in part explained by similar brain architectures, a large proportion is likely driven by task engagement. In contrast, ISFC was not significantly higher on any days of learning (mean ISFC during rest was 0.73, *p* > 0.1). Finally, it is important to note that we observed a significant correlation between each subject’s global ISFC during learning and their global ISFC during the resting state (*r*=0.97, *p*=2.71 × 10^−13^).

In sum, on day one of learning, brain activity is significantly more constrained than during rest, while there is a slight but not significant decrease in ISFC from rest to task. On day two, both ISFC and ISC increase significantly. Later in learning (days three and four), ISFC remains high, while ISC decreases. Given that subjects’ ISFC was strongly correlated between rest and task, much of the ISFC is likely driven by constraints of brain architecture independent of the task. However, these dynamics of high ISC on day one and high ISFC on day two point to a potential driver-follower mechanism of constrained activity preceding constrained functional connectivity over the four days of value learning, with this increase in ISFC likely being caused, in part, by task engagement.

### Regional variability of functional constraints during learning

If the dynamics of whole-brain inter-subject connectivity map on to different phases of value learning, it is natural to ask which brain systems might be most complicit in these phases. To address this question, we first partitioned brain regions into objectively defined functional modules using the GenLouvain community detection algorithm [49] (see Materials and Methods). Briefly, community detection is applied to the functional network constructed from data obtained during each task session. This procedure parses brain regions into functional modules such that brain regions within the same module exhibit strong functional connections, and brain regions between different modules exhibit weak functional connections. To obtain a single representative partition of brain regions into modules across subjects and scans, we computed the module allegiance matrix [39], which encodes the probability that any two regions belong to the same functional module across all data (Fig 5a). By applying a final round of community detection to the module allegiance matrix [38], we identified seven modules that were associated with different putative brain systems, including a fronto-temporal module (FT) which covered most of the limbic lobe, a sensorimotor module (SM), an auditory module (AUD) including hippocampus and amygdala, the common default mode system (DMN), a language & memory module (LAN), a visual module (VIS), and a subcortical module composed of the putamen, caudate, and thalamus (PCT) (Fig. 5b). We found that the ISFC was significantly higher in the DMN, SM, and VIS modules than in the remaining module (*p* < 0.05; Fig. 5c).

**Figure 5.**
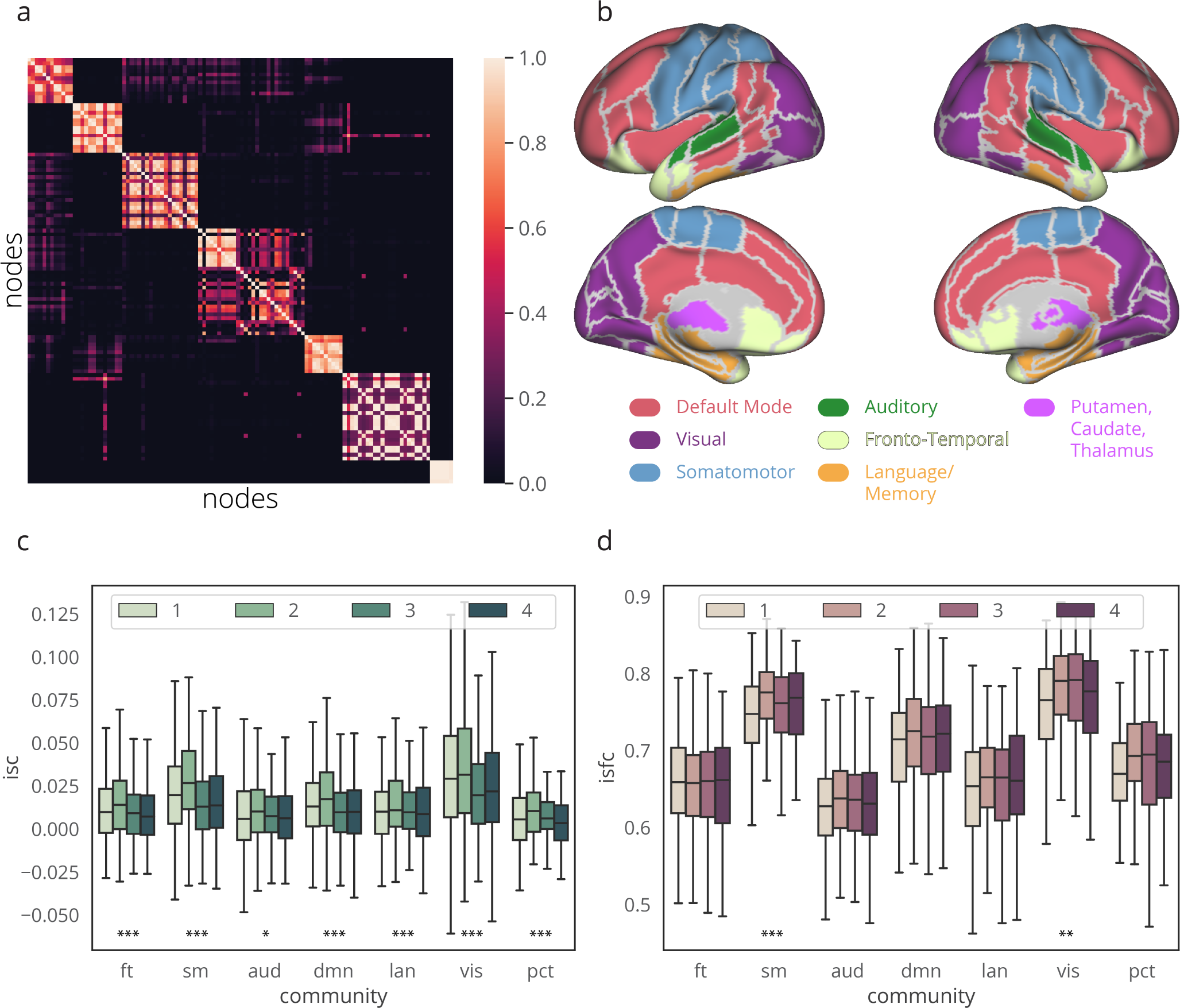
Variation in ISC and ISFC across functional modules. **a** The module allegiance matrix, each element of which indicates the probability that two brain regions belong to the same community across all functional networks estimated from fMRI data acquired during task performance. Using a common community detection method, we identified seven communities across all scanning sessions for all of the subjects. **b** We projected the seven communities onto the MRI surface (fslr32k), and we named each of them based on their anatomical locations: frontal temporal module (FT), sensorimotor module (SM), auditory module (AUD), default mode system (DMN), language module (LAN), visual module (VIS), and a subcortical module composed of the putamen, caudate, and thalamus (PCT). **c**–**d** In general, we observed that the ISC and ISFC for each network module displayed the same day-to-day pattern as the whole-brain average ISC and ISFC. Using a repeated measures ANOVA, we determined whether there was a statistically significant effect of day for each of the modules. Statistical significance for the ANOVA with the measured effect across the four days are reported as follows: \**p* < 0.05; \*\**p* < 0.01; \*\*\**p* < 0.001.

Next, we assessed the degree to which the time-dependent variation in ISC and ISFC might differ across the seven brain modules (Fig. 6). With the exception of the FT module’s ISFC, all of the modules display similar profiles of ISFC and ISC as shown for the whole brain in Fig. 4c), with each module showing a significant difference in ISC and ISFC between days one and two (*t* > 2.0, *p* > 0.05). ISC was initially higher than ISFC on day one, and then ISFC became higher than ISC on day two. Finally, both ISC and ISFC decrease on days three and four, and ISFC remains equal to or higher than ISC. The distinct temporal signature of the FT module may be due to the fact that these regions have been associate with the learning of value and thus strong activity, but not necessarily strong ISFC, in these regions might dominate early learning. Taken together, this pattern of findings implies that the temporal evolution of inter-subject connectivity and activity is similar across the entire brain.

**Figure 6.**
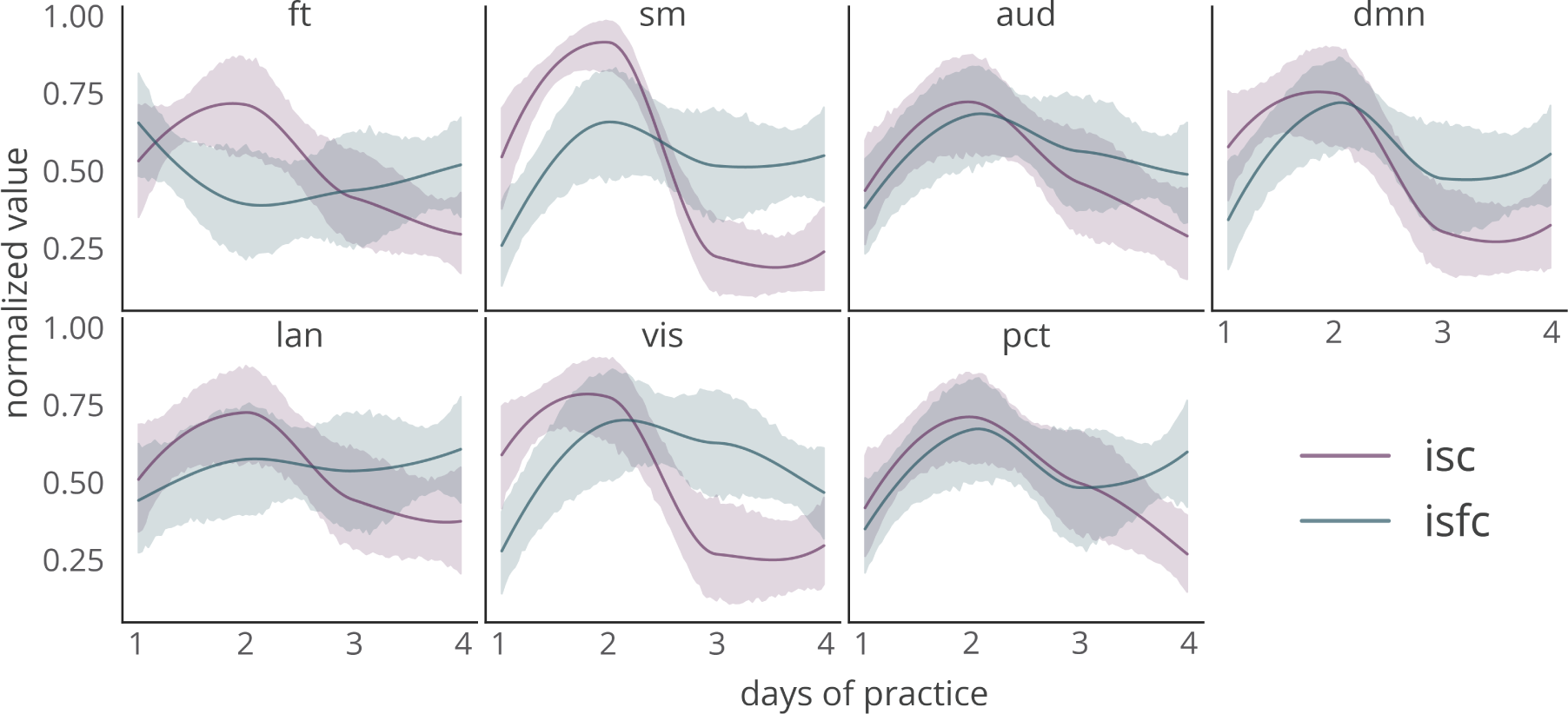
Module-specific time-dependent variation in ISC and ISFC. The normalized ISC (red) and ISFC (blue) curves for the seven network modules, ordered from greatest to least difference between ISC and ISFC in the shape of the curve. Broadly, we observed a significant difference between normalized ISC and normalized ISFC on day 3 and day 4, particularly for the SM, VIS, DMN and AUD modules. These differences occurred as ISC significantly decreased with training while ISFC remained high.

### Relation between ISFC and ISC and individual differences in value learning

Lastly, we asked whether inter-individual differences in the constraints of activity and functional connectivity could explain inter-individual differences in learning. We observed that average task accuracy increased across the four days of task practice (Fig. 7a; two-way ANOVA: *F*(3,79) = 3.69, *p* = 0.0169). Next, we sought to determine whether the extent to which a subject’s activity or connectivity was constrained to the group level was indicative of learning performance. To address this question, we used a linear regression model to fit each community’s ISC or ISFC in each subject to learning performance. The model was fit to all subjects and all days except one subject on one day, and the model was then used to predict the left out subject’s performance on that day. Notably, the model was only able to significantly relate ISFC, not ISC, to learning performance, suggesting that learning performance is related to constraints on connectivity but not activity (Fig. 7b).

**Figure 7.**
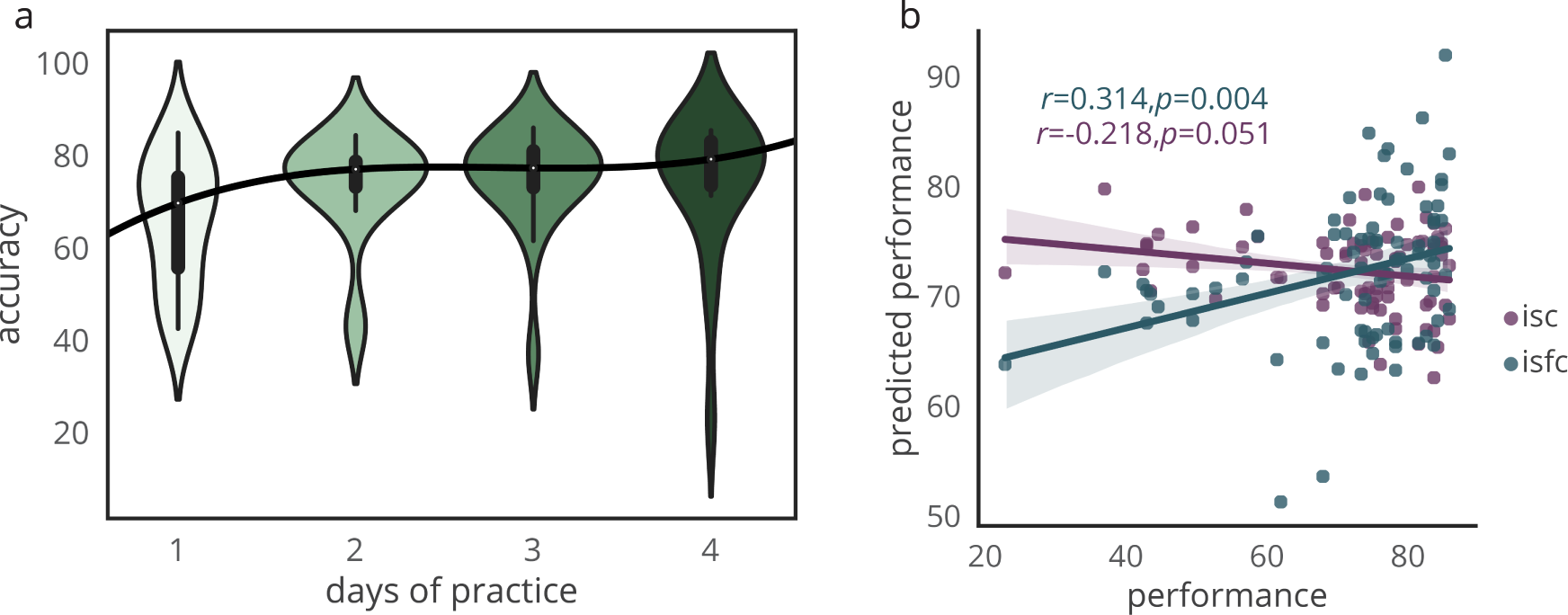
The relation between inter-individual differences in ISC and ISFC, and individual differences in task performance. **a**, Accuracy across subjects increased with days of practice. **b**, A linear regression model was fit to relate each community’s ISC or ISFC in each subject to learning performance. The model was fit to all subjects and all days except one subject on one day, and the model was then used to predict the left out subject’s performance on that day. The model was only able to significantly relate ISFC, not ISC, to task performance.

## Discussion

In this study, we used inter-subject analyses to determine the relative roles of activity and connectivity constraints during the learning of object values over the course of four days of learning. Importantly, our analyses were able to separate evidence for group constraints on large-scale neuronal processes supporting learning from evidence for idiosyncratic processes that may be relevant for task performance in single subjects. In general, our results suggest that connectivity and activity tend to be constrained to the group average in a similar manner across brain regions, but in a distinct manner across time. Greater inter-subject correlations in both activity and connectivity exist in early versus late learning, suggesting that the early stages of learning recruit common neurophysiological processes necessary for encoding information. We observed greater inter-subject correlations in activity than functional connectivity on day one, likely because stimulus induced activity is very similar across individuals. However, we observed greater inter-subject correlations in functional connectivity than in activity late in learning, suggesting a role of connectivity in the consolidation of learned information, and greater individual variation in activity patterns in late learning that might serve to optimize behavior in an idiosyncratic manner. Collectively, our findings demonstrate that subjects express common stimulus-induced activity and connectivity that evolves based on the stage of the learning process.

### Module specific connectivity constraints during value learning

Are some cognitive systems more constrained in their activity and connectivity than others? We identified several data-derived network modules reminiscent of cognitive systems – visual, sensory-motor, and default mode modules – that demonstrated significantly greater intersubject activity and intersubject functional connectivity than other modules. In particular, we observed significantly higher ISC and ISFC within lateral occipital cortex, lingual gyrus, fusiform gyrus, and pericalcarine cortex, as well as other areas of the occipital lobe, suggesting that value learning commonly recruits brain regions responsible for visual perception, visual memory, and decision-making based on chosen value [50]. We also observed significant ISC and ISFC in temporal and parietal areas involved in attention, mentalizing, and the attribution of self-belief [51, 52]. It is intuitively plausible that the significant ISC and ISFC in visual and sensorimotor systems is a direct result of shared sensory inputs. However, the significant ISC and ISFC in the default mode system is unlikely to be adequately explained by the same mechanism, due to the tendency of this system to be active independent from stimulus input [53]. An alternative explanation lies in the fact that default mode areas typically display high degree in network models, comprising a large part of the rich club [54], supporting the notion that the default mode forms a stable core of areas that exhibit similar activity and connectivity across subjects.

Interestingly, we found that the regions for which their ISC was high and also associated with value were primarily in orbital frontal cortex, while the regions for which their ISFC was high and also associated with value were primarily in the canonical fronto-parietal network [48]. Additionally, we observed that several brain regions displayed ISFC that exceeded their ISC, including the amygdala, thalamus, precuneus, cuneus, cingulate, and lateral frontal cortex. Collectively, these regions are commonly associated with higher order cognitive functions such as flexibility of thinking, problem solving, cognitive inhibition, and attentional control [51]. One explanation for the high ISFC exhibited by these regions is that they represent important cognitive control areas that are likely to act as functional hubs that may be commonly integrated with other brain regions across subjects [55, 54, 56]. Specifically, frontal pole frequently interacts with the anterior cingulate gyrus during reward-guided learning [57], and the precuneus – although widely known to serve as an important hub in the default mode [58] – is often activated during episodic memory retrieval and self-processing operations [59]. The higher ISFC at the precuneus might be related to the higher reaction times in correct value trials observed during value learning tasks [60], which is thought to be driven in part by a subject’s memory of similar tasks experienced previously. In summary, more constrained brain activity appears to occur within predominantly sensory-specific brain regions, and more constrained functional connectivity appears to occur in integrative areas that are commonly associated with higher order cognitive function.

### Time-dependent variation in constraints during learning

An advantage of our experimental approach is the ability to examine dynamic changes in neuronal processes, and their variation across subjects, throughout different phases of value learning. During the early stages of learning, we observed marked constraints on activity, suggesting that participants generally exhibited a similar time course of BOLD activation in response to task demands. During later stages of learning, we observed a decrease in constraints on activity. We speculate that this shift from high to low constraints might reflect individualized patterns of activity that support inter-individual variability in learning strategies as well as the fact that in later stages of learning greater neural real estate is available for contemporaneous, non-task-related processing. In contrast, constraints on functional connectivity gradually increased between the first and second day, and then subsequently decreased in day 3, thereafter remaining relatively constant on day 4. This empirically observed trajectory of functional connectivity constraints is reminiscent of the typical shape of a traditional learning curve [61], suggesting that functional connectivity may comprise a general adaptive mechanism for learning [62, 63, 64].

Importantly, we found that putative cognitive systems displayed differential patterns of learning-induced ISFC and ISC. The high ratio of ISFC-to-ISC observed in the sensorimotor system could be attributed to common inputs to the sensory system for the processing of stimulus information. The complimentary increase in the ISFC of the visual and sensorimotor systems may result from trained motor coordination of hand and finger movements to press the button for task response [37, 65]. The resulting changes suggest a role in visual and sensorimotor systems in supporting visual identification of objects, interpretation of value, and motor coordination of hand-finger movement to guide more accurate and efficient decision making after the critical learning period. The lower ISC in the later stage of learning could be ascribed to higher individual differences in time courses of BOLD signals after adaptation to the task and once the procedure is more thoroughly learned [66, 67]. Interestingly, we find differential involvement of the default mode over the course of value learning: connectivity within the default mode system becomes more generalized across subjects, and activity within the default mode system becomes more individualized across subjects. Although the default mode system is often viewed as a task-negative system that is typically more active during resting state processes, here we report evidence that the architecture of this system is highly conserved across learning.

### Functional network drivers of value learning

Finally, it is important to consider the question of whether, and to what degree, high ISC and ISFC might support or hamper learning. We find that ISFC, but not ISC, was significantly correlated with task accuracy, suggesting that behavior might be improved when the functional network reorganizes according to subject-general constraints on neural processes, but that stimulus-induced activity and learning-related activity constraints are not relevant to learning performance. While individual variation in stimulus-induced activity might be responsible for modulating subject-specific behavior during value learning tasks [68, 69, 70], the extent to which the activity deviated from the group level constraints did not hamper or help performance. The role of ISFC in predicting individual differences in performance is consistent with prior work uncovering relations between task performance and patterns of functional connectivity motor-visual learning [71] and reinforcement learning [72]. It will be of interest in future to develop experimental paradigms that can separately probe the performance-relevant role of dynamic functional connectivity in the motor visual system, in the reward system, and in areas related to object perception and object value, all within the same cohort of health subjects. Moreover, it would be of interest to test whether these roles diverge in patients with deficits in motor-visual function or reward processing.

### Conclusions

Our study offers initial evidence that brain activity and functional connectivity display specific regional and temporal constraints across individuals during distinct stages of learning the value of novel objects. Activity and connectivity are most constrained in motor and visual regions, as well as regions associated with value. Activity is most similar across subjects on the first day of learning the task, while the similarity of functional connectivity peaks on the second day when the task itself is well understood, but the behavior is still adapting swiftly. While constraints on both activity and functional connectivity decrease in the later stages of learning, functional connectivity remains more constrained than activity. Notably, how constrained an individual’s connectivity is to the group level is predictive of individual differences in task accuracy. Collectively, our finding pave the way for concerted efforts to better understand the relation between activity and connectivity as two important and distinct phenotypes of human learning, and the relative task-induced constraints that impinge on each.

## Acknowledgements

D.S.B. acknowledges support from the John D. and Catherine T. MacArthur Foundation, the Alfred P. Sloan Foundation, the ISI Foundation, the Paul G. Allen Foundation, the Army Research Laboratory (W911NF-10-2-0022), the Army Research Office (Bassett-W911NF-14-1-0679, Grafton-W911NF-16-1-0474, DCIST-W911NF-17-2-0181), the Office of Naval Research, the National Institute of Mental Health (2-R01-DC009209-11, R0-MH112847, R01-MH107235, R21-MH106799, R01-MH113550), the National Institute of Child Health and Human Development (1R01-HD086888-01), National Institute of Neurological Disorders and Stroke (R01-NS099348), and the National Science Foundation (BCS-1441502, BCS-1430087, NSF PHY-1554488 and BCS-1631550). The content is solely the responsibility of the authors and does not necessarily represent the official views of any of the funding agencies.

## References

[1] E. S. Finn, X. Shen, D. Scheinost, M. D. Rosenberg, J. Huang, M. M. Chun, X. Papademetris, R. T. Constable, Functional connectome fingerprinting: identifying individuals using patterns of brain connectivity, Nature Neuroscience 18 (11) (2015) 1664–1671, doi:10.1038/nn.4135, URL https://doi.org/10.1038%2Fnn.4135.

[2] M. A. Bertolero, B. T. T. Yeo, D. S. Bassett, M. D’Esposito, A mechanistic model of connector hubs, modularity and cognition, Nature Human Behaviour 2 (10) (2018) 765–777.

[3] D. S. Bassett, M. Yang, N. F. Wymbs, S. T. Grafton, Learning-induced autonomy of sensorimotor systems, Nature neuroscience 18 (5) (2015) 744–751.

[4] M. G. Mattar, N. F. Wymbs, A. S. Bock, G. K. Aguirre, S. T. Grafton, D. S. Bassett, Predicting future learning from baseline network architecture, Neuroimage 172 (2018) 107–117.

[5] G. Schoenbaum, A. A. Chiba, M. Gallagher, Changes in functional connectivity in orbitofrontal cortex and basolateral amygdala during learning and reversal training, Journal of Neuroscience 20 (13) (2000) 5179–5189.

[6] E. M. Gordon, T. O. Laumann, A. W. Gilmore, D. J. Newbold, D. J. Greene, J. J. Berg, M. Ortega, C. Hoyt-Drazen, C. Gratton, H. Sun, J. M. Hampton, R. S. Coalson, A. L. Nguyen, K. B. McDermott, J. S. Shimony, A. Z. Snyder, B. L. Schlaggar, S. E. Petersen, S. M. Nelson, N. U. F. Dosenbach, Precision Functional Mapping of Individual Human Brains, Neuron 95 (4) (2017) 1–35.

[7] I. Tavor, O. P. Jones, R. B. Mars, S. M. Smith, T. E. Behrens, S. Jbabdi, Task-free MRI predicts individual differences in brain activity during task performance, Science 352 (6282) (2016) 216–220.

[8] R. F. Betzel, D. S. Bassett, Multi-scale brain networks, Neuroimage.

[9] C. Gratton, T. O. Laumann, A. N. Nielsen, D. J. Greene, E. M. Gordon, A. W. Gilmore, S. M. Nelson, R. S. Coalson, A. Z. Snyder, B. L. Schlaggar, N. U. F. Dosenbach, S. E. Petersen, Functional Brain Networks Are Dominated by Stable Group and Individual Factors, Not Cognitive or Daily Variation, Neuron 98 (2) (2018) 1–20.

[10] M. L. Seghier, C. J. Price, Visualising inter-subject variability in fMRI using threshold-weighted overlap maps, Scientific Reports 6 (2016) 20170 EP –.

[11] D. S. Bassett, M. G. Mattar, A network neuroscience of human learning: Potential to inform quantitative theories of brain and behavior, Trends in Cognitive Sciences.

[12] A. DiMartino, D.A. Fair, C. Kelly, T. D. Satterthwaite, F. X. Castellanos, M. E. Thomason, R. C. Craddock, B. Luna, B. L. Leventhal, X.-N. Zuo, et al., Unraveling the miswired connectome: a developmental perspective, Neuron 83 (6) (2014) 1335–1353.

[13] O. Atun-Einy, S. E. Berger, A. Scher, Pulling to stand: common trajectories and individual differences in development., Developmental psychobiology 54 (2) (2012) 187–198, ISSN 1098–2302, doi:10.1002/dev.20593, URL http://www.ncbi.nlm.nih.gov/pubmed/21815138.

[14] R. Harrison, Learning and development, Development and Learning in Organizations: An International Journal 26 (1).

[15] A. Wigfield, J. S. Eccles, Expectancy–value theory of achievement motivation, Contemporary educational psychology 25 (1) (2000) 68–81.

[16] A. M. Dale, B. Fischl, M. I. Sereno, Cortical surface-based analysis: I. Segmentation and surface reconstruction, Neuroimage 9 (2) (1999) 179–194.

[17] D. N. Greve, B. Fischl, Accurate and robust brain image alignment using boundary-based registration, Neuroimage 48 (1) (2009) 63–72.

[18] M. Jenkinson, Improving the registration of B0-disorted EPI images using calculated cost function weights, Neuroimage 22 (2004) e1544–e1545.

[19] M. Jenkinson, P. Bannister, M. Brady, S. Smith, Improved optimization for the robust and accurate linear registration and motion correction of brain images, Neuroimage 17 (2) (2002) 825–841.

[20] S. M. Smith, Fast robust automated brain extraction, Human brain mapping 17 (3) (2002) 143–155.

[21] K. J. Friston, S. Williams, R. Howard, R. S. Frackowiak, R. Turner, Movement-related effects in fMRI time-series, Magnetic resonance in medicine 35 (3) (1996) 346–355.

[22] Y. Behzadi, K. Restom, J. Liau, T. T. Liu, A component based noise correction method (CompCor) for BOLD and perfusion based fMRI, Neuroimage 37 (1) (2007) 90–101.

[23] H. J. Jo, Z. S. Saad, W. K. Simmons, L. A. Milbury, R. W. Cox, Mapping sources of correlation in resting state FMRI, with artifact detection and removal, Neuroimage 52 (2) (2010) 571–582.

[24] X. J. Chai, A. N. Castañón, D. Öngür, S. Whitfield-Gabrieli, Anticorrelations in resting state networks without global signal regression, Neuroimage 59 (2) (2012) 1420–1428.

[25] K. Murphy, R. M. Birn, D. A. Handwerker, T. B. Jones, P. A. Bandettini, The impact of global signal regression on resting state correlations: are anti-correlated networks introduced?, Neuroimage 44 (3) (2009) 893–905.

[26] S. M. Smith, M. Jenkinson, M. W. Woolrich, C. F. Beckmann, T. E. J. Behrens, H. Johansen-Berg, P. R. Bannister, M. De Luca, I. Drobnjak, D. E. Flitney, R. K. Niazy, J. Saunders, J. Vickers, Y. Zhang, N. De Stefano, J. M. Brady, P. M. Matthews, Advances in Functional and Structural MR Image Analysis and Implementation as FSL, Neuroimage 23 (2004) S208–S219.

[27] M. W. Woolrich, S. Jbabdi, B. Patenaude, M. Chappell, S. Makni, T. E. J. Behrens, C. Beckmann, M. Jenkinson, S. M. Smith, Bayesian Analysis of Neuroimaging Data in FSL, Neuroimage 45 (1) (2009) S173–S186.

[28] C. T. Butts, Revisiting the foundations of network analysis, Science 325 (5939) (2009) 414–416.

[29] D. S. Bassett, P. Zurn, J. I. Gold, On the nature and use of models in network neuroscience, Nat Rev Neurosci Epub ahead of print.

[30] C. Torrence, G. P. Compo, A Practical Guide to Wavelet Analysis, Bulletin of the American Meteorological Society 79 (1) (1998) 61–78.

[31] B. Cazelles, M. Chavez, G. C. de Magny, J.-F. Guegan, S. Hales, Time-Dependent Spectral Analysis of Epidemiological Time-Series with Wavelets, Journal of the Royal Society Interface 4 (15) (2007) 625–636.

[32] A. Grinsted, J. C. Moore, S. Jevrejeva, Application of the Cross Wavelet Transform and Wavelet Coherence to Geophysical Time Series, Nonlinear Processes in Geophysics 11 (5/6) (2004) 561–566.

[33] O. Sporns, R. F. Betzel, Modular Brain Networks, Annu Rev Psychol 67 (2016) 613–640.

[34] M. Girvan, M. E. J. Newman, Community Structure in Social and Biological Networks, Proc. Natl. Acad. Sci. USA 99 (2001) 8271–8276.

[35] P. De Meo, E. Ferrara, G. Fiumara,, A. Provetti, Generalized Louvain Method for Community Detection in Large Networks, Intelligent Systems Design and Applications (2011) 88–93.

[36] P. J. Mucha, T. Richardson, K. Macon, M. A. Porter, J.-P. Onnela, Community Structure in Time-Dependent, Multiscale, and Multiplex Networks, Science 328 (5980) (2010) 876–878.

[37] D. S. Bassett, N. F. Wymbs, M. A. Porter, P. J. Mucha, J. M. Carlson, S. T. Grafton, Dynamic reconfiguration of human brain networks during learning, Proceedings of the National Academy of Sciences 108 (18) (2011) 7641–7646.

[38] D. Bassett, M. Porter, N. Wymbs, S. Grafton, J. Carlson, P. Mucha, Robust detection of dynamic community structure in networks., Chaos 23 (2013) 013142.

[39] D. Bassett, M. Yang, N. Wymbs, S. Grafton, Learning-induced autonomy of sensorimotor systems., Nat Neurosci 18 (2015) 744–751.

[40] B. F. Manly, Randomization, bootstrap and Monte Carlo methods in biology, vol. 70, CRC Press, 2006.

[41] D. A. Handwerker, V. Roopchansingh, J. Gonzalez-Castillo, P. A. Bandettini, Periodic changes in fMRI connectivity, Neuroimage 63 (3) (2012) 1712–1719.

[42] E. D. Van, S. Smith, D. Barch, T. Behrens, E. Yacoub, K. Ugurbil, The WU-Minn Human Connectome Project: an overview., Neuroimage 80 (2013) 62–79.

[43] A. Schaefer, R. Kong, E. Gordon, T. Laumann, X. Zuo, A. Holmes, S. Eickhoff, B. Yeo, Local-Global Parcellation of the Human Cerebral Cortex from Intrinsic Functional Connectivity MRI., Cereb Cortex (2017) 1–20.

[44] M. G. Mattar, S. L. Thompson-Schill, D. S. Bassett, The network architecture of value learning, Network Neuroscience.

[45] A. S. Persichetti, G. K. Aguirre, S. L. Thompson-Schill, Value is in the eye of the beholder: early visual cortex codes monetary value of objects during a diverted attention task, Journal of cognitive neuroscience.

[46] A. W. Kemp, B. F. J. Manly, Randomization Bootstrap and Monte Carlo Methods in Biology., Biometrics 53 (4) (1997) 1560, doi:10.2307/2533527, URL https://doi.org/10.2307%2F2533527.

[47] T. Yarkoni, R. A. Poldrack, T. E. Nichols, D. C. Van Essen Nature, 2011, Large-scale automated synthesis of human functional neuroimaging data, nature.com.

[48] B. T. Thomas Yeo, F. M. Krienen, J. Sepulcre, M. R. Sabuncu, D. Lashkari, M. Hollinshead, J. L. Roffman, J. W. Smoller, L. Zöllei, J. R. Polimeni, B. Fischl, H. Liu, R. L. Buckner, The organization of the human cerebral cortex estimated by intrinsic functional connectivity, Journal of Neurophysiology 106 (3) (2011) 1125–1165.

[49] P. De Meo, E. Ferrara, G. Fiumara, A. Provetti, Generalized louvain method for community detection in large networks, in: Intelligent Systems Design and Applications (ISDA), 2011 11th International Conference on, IEEE, 88–93, 2011.

[50] S.-G. Kuai, D. Levi, Z. Kourtzi, Learning optimizes decision templates in the human visual cortex, Current Biology 23 (18) (2013) 1799–1804.

[51] M. Corbetta, G. L. Shulman, Control of goal-directed and stimulus-driven attention in the brain, Nature reviews. Neuroscience 3 (3) (2002) 201.

[52] M. Corbetta, G. L. Shulman, Spatial neglect and attention networks, Annual review of neuroscience 34 (2011) 569–599.

[53] D. S. Margulies, S. S. Ghosh, A. Goulas, M. Falkiewicz, J. M. Huntenburg, G. Langs, G. Bezgin, S. B. Eickhoff, F. X. Castellanos, M. Petrides, E. Jefferies, J. Smallwood, Situating the default-mode network along a principal gradient of macroscale cortical organization, Proceedings of the National Academy of Sciences 113 (44) (2016) 12574–12579, doi:10.1073/pnas.1608282113, URL https://doi.org/10.1073%2Fpnas.1608282113.

[54] M. A. Bertolero, B. T. T. Yeo, M. D’Esposito, The diverse club, Nature Communications 8 (1) (2017) 1277, doi:10.1038/s41467-017-01189-w.

[55] K. Hwang, M. A. Bertolero, W. B. Liu, M. D’Esposito, The Human Thalamus Is an Integrative Hub for Functional Brain Networks., The Journal of neuroscience: the official journal of the Society for Neuroscience 37 (23) (2017) 5594–5607.

[56] M. A. Bertolero, B. T. T. Yeo, M. D’Esposito, The modular and integrative functional architecture of the human brain, Proceedings of the National Academy of Sciences 112 (49) (2015) E6798–E6807, doi:10.1073/pnas.1510619112, URL https://doi.org/10.1073%2Fpnas.1510619112.

[57] M. F. Rushworth, M. P. Noonan, E. D. Boorman, M. E. Walton, T. E. Behrens, Frontal cortex and reward-guided learning and decision-making, Neuron 70 (6) (2011) 1054–1069.

[58] M. E. Raichle, A. M. MacLeod, A. Z. Snyder, W. J. Powers, D. A. Gusnard, G. L. Shulman, A default mode of brain function, Proceedings of the National Academy of Sciences 98 (2) (2001) 676–682.

[59] A. E. Cavanna, M. R. Trimble, The precuneus: a review of its functional anatomy and behavioural correlates, Brain 129 (3) (2006) 564–583.

[60] K. Oishi, K. Toma, E. T. Bagarinao, K. Matsuo, T. Nakai, K. Chihara, H. Fukuyama, Activation of the precuneus is related to reduced reaction time in serial reaction time tasks, Neuroscience research 52 (1) (2005) 37–45.

[61] R. A. Rescorla, Variation in the effectiveness of reinforcement and nonreinforcement following prior inhibitory conditioning, Learning and motivation 2 (2) (1971) 113–123.

[62] E. H. Baeg, Y. B. Kim, J. Kim, J.-W. Ghim, J. J. Kim, M. W. Jung, Learning-induced enduring changes in functional connectivity among prefrontal cortical neurons, Journal of Neuroscience 27 (4) (2007) 909–918.

[63] Z. Fatima, N. Kovacevic, B. Misic, A. R. McIntosh, Dynamic functional connectivity shapes individual differences in associative learning, Human brain mapping 37 (11) (2016) 3911–3928.

[64] R. Patel, R. N. Spreng, G. R. Turner, Functional brain changes following cognitive and motor skills training: a quantitative meta-analysis, Neurorehabilitation and neural repair 27 (3) (2013) 187–199.

[65] I. Toni, N. Ramnani, O. Josephs, J. Ashburner, R. E. Passingham, Learning arbitrary visuomotor associations: temporal dynamic of brain activity, Neuroimage 14 (5) (2001) 1048–1057.

[66] A. J. Bastian, Understanding sensorimotor adaptation and learning for rehabilitation, Current opinion in neurology 21 (6) (2008) 628.

[67] A. J. Keller, R. Houlton, B. M. Kampa, N. A. Lesica, T. D. Mrsic-Flogel, G. B. Keller, F. Helmchen, Stimulus relevance modulates contrast adaptation in visual cortex, eLife 6 (2017) e21589.

[68] R. T. Gerraty, J. Y. Davidow, G. E. Wimmer, I. Kahn, D. Shohamy, Transfer of learning relates to intrinsic connectivity between hippocampus, ventromedial prefrontal cortex, and large-scale networks, Journal of Neuroscience 34 (34) (2014) 11297–11303.

[69] A. R. Laird, P. M. Fox, S. B. Eickhoff, J. A. Turner, K. L. Ray, D. R. McKay, D. C. Glahn, C. F. Beckmann, S. M. Smith, P. T. Fox, Behavioral interpretations of intrinsic connectivity networks, Journal of cognitive neuroscience 23 (12) (2011) 4022–4037.

[70] M. Yamashita, M. Kawato, H. Imamizu, Predicting learning plateau of working memory from whole-brain intrinsic network connectivity patterns, Scientific reports 5 (2015) 7622.

[71] M. H. Heitger, R. Ronsse, T. Dhollander, P. Dupont, K. Caeyenberghs, S. P. Swinnen, Motor learning-induced changes in functional brain connectivity as revealed by means of graph-theoretical network analysis, Neuroimage 61 (3) (2012) 633–650.

[72] R. T. Gerraty, J. Y. Davidow, K. Foerde, A. Galvan, D. S. Bassett, D. Shohamy, Dynamic flexibility in striatal-cortical circuits supports reinforcement learning, J Neurosci (2018) 2084–2017.

